# Myosin VI controls localization of Golgi satellites at active presynaptic boutons

**DOI:** 10.1101/2024.05.08.593268

**Authors:** Nathalie Hertrich, Nathanael Doil, Anja Konietzny, Marina Mikhaylova

**Author notes:** **Conflict of interest:** The authors declare no competing financial interests.

## Abstract

Neurons, as long-living non-dividing cells with complex morphology, depend on highly elaborate secretory trafficking system which ensures the constant delivery, removal and recycling of proteins and membranes. Previously, we have shown that simplified Golgi-related structures called Golgi satellites (GS), distinct from the somatic Golgi complex, are present in dendrites of primary hippocampal neurons and are involved in glycosylation and local forward trafficking of membrane proteins. However, whether GS are also targeted to axons of principal neurons have not been explored. Here, we investigate the subcellular distribution of GS in adult hippocampal neurons and discover that mobile and stationary GS are present along the entire axonal length, extending to the distal tips of the growth cone. Live imaging experiments revealed that neuronal firing modulates the switch between long range transport mediated by kinesin and dynein and stalling. We found that GS frequently pause or stop at pre-synaptic sites in activity-dependent manner. This behavior depends on the actin cytoskeleton and the actin-based motor protein myosin VI. Overall, our study demonstrates that neuronal activity can dynamically regulate the positioning of GS in the axon, shedding light on the intricate mechanisms underlying organelle trafficking in neurons.

**Significance statement:** Our study unveils the presence and dynamic behavior of Golgi satellites (GS), specialized organelles implicated in glycosylation and membrane protein trafficking, within axons of adult neurons. We found that mobile and stationary GS are present throughout the axonal length, including distal growth cone regions. GS are transported bidirectionally and their preferential pausing at presynaptic sites is regulated by neuronal firing. GS positioning at presynaptic boutons relies on the actin cytoskeleton and the myosin VI motor protein. This work elucidates how neuronal activity modulates GS distribution, shedding light on fundamental mechanisms of organelle trafficking in neurons.

## Introduction

Neurons are highly polarized cells with distinct axonal and somatodendritic domains separated by the pre-axonal exclusion zone and the axon initial segment. As long-living non-dividing cells with a complex morphology, neurons depend on the constant delivery, removal and recycling of proteins and membranes. To accommodate these demands, they developed an elaborate and well-controlled secretory trafficking system which spans the entire neuron, reaching even the distal tips of axons and dendrites. Secretory organelles such as the endoplasmic reticulum (ER), Golgi-related compartments, various types of endosomes, autophagosomes, and lysosomes are found in the soma as well as in the dendrite and axon. Their transport and correct localization are essential for neuronal survival. If impaired, this may lead to severe diseases such as lysosomal storage diseases, Alzheimer’s disease and Parkinson’s disease (Wang et al., 2014; Pará et al., 2020).

The Golgi apparatus, consisting of flattened tubules called cisternae, plays a central role in modifying, sorting, and packaging proteins and lipids into vesicles for transport. Proteins and lipids pass through the Golgi complex, entering from its cis-face and exiting through its trans-face. Along the way, they undergo various post-translational modifications such as glycosylation, phosphorylation, removal of mannose residues, and others. In addition to the somatic Golgi complex, stacked tubular Golgi structures termed Golgi outposts (GO) have been described at the branch points of Drosophila neurons and in the apical dendrite of excitatory pyramidal neurons in rodents (Hanus and Ehlers, 2008; Ori-McKenney et al., 2012). GO are frequently clustered near ER exit sites (ERES), and are involved in various local functions such as protein secretion, trafficking, and glycosylation (Horton et al., 2005; Wang et al., 2012; Kemal et al., 2022). Previous studies have revealed another Golgi-related organelle, Golgi satellites (GS) (Mikhaylova et al., 2016), which are widely distributed in all dendrites of primary hippocampal neurons. In contrast to GO, GS have a more simplified structure. They do not contain cis-Golgi elements and do not form membrane stacks (Andres-Alonso et al., 2023). Similar to GOs, GS contain a glycosylation machinery and are located in close proximity to the ER and ERGIC, forming a local secretory system where proteins can be recycled, and newly synthesized ER proteins can pass through. A recent study has shown that GS are essential for the polysialylation of neural cell adhesion molecules (NCAM) in distal dendrites (Andres-Alonso et al., 2023). In addition, GS are involved in shaping the dendritic surface glycoproteome of neurons (Govind et al., 2021).

In axons, Golgi-related compartments are not well investigated with only a few descriptive studies reporting on them. In the peripheral nervous system, Golgi components have been reported to cluster in axons of the rat sciatic nerve, and GS have been described to be involved in trafficking of cold-sensitive TRPM8 ion channels in axons of dorsal-root ganglion (DRG) neurons (González et al., 2016; Cornejo et al., 2020). In the central nervous system, while trans-Golgi carriers have been shown to deliver lysosomal proteases (cathepsins) from the soma to the distal axon of cortical neurons, much less is known about the distribution and trafficking of GS (Lie et al., 2021). It is still an open question whether GS exist in axons and how they are transported and distributed.

Long-range transport of organelles between the soma, dendrites, and axon is mediated by the microtubule (MT) cytoskeleton and kinesin and dynein motor proteins, whereas short-range transport and anchoring at specific sites rely on the F-actin cytoskeleton and the myosin motor protein family. Although not previously explored, it is possible that GS undergo MT and F-actin-based transport in both dendrites and axons of mammalian neurons.

Here, we investigated the distribution and trafficking of GS in axons of adult rat hippocampal neurons. We found that GS are present along the entire axonal length, including growth cones. Their bidirectional motility and the switch between long-range transport and anchoring at specific sites is regulated by neuronal activity and myosin VI motor protein.

## Materials and Methods

### Animals

Wistar Unilever HsdCpb:WU (Envigo) rats were obtained from the animal facility of the University Medical Center Hamburg-Eppendorf, UKE, Hamburg, Germany, from the animal facility of the Humboldt-Universität zu Berlin and from Janvier laboratories. Protocols for prepping primary neuronal cultures were approved by the local authorities of the city-state Hamburg (Behörde für Gesundheit und Verbraucherschutz, Fachbereich Veterinärwesen), the animal care committee of the University Medical Center Hamburg-Eppendorf as well as the local authorities of the city-state Berlin (Landesamt für Gesundheit und Soziales).

### Key resources, reagents, constructs and cloning

A detailed information of the antibodies, DNA plasmids and pharmacological compounds used in this study can be found in the Appendix Table S1. All constructs used in this study were verified by sequencing and are summarized in the Appendix Table S1. Restriction enzymes were purchased from Thermo Fisher Scientific. The St3gal5-GFP plasmid was cloned by PCR amplification of the St3gal5-GFP (Mikhaylova et al., 2016) and inserting it into into a pAAV backbone using EcoRI and HindIII restriction sites. The pORANGE_Actin was produced by exchanging the GFP in the Addgene #139666 vector (pORANGE_Actin#2-GFP) for a TagRFP (amplified from pTagRFP-C, Evrogen) using the restriction enzymes NheI and BamHI. The St3gal5-Halo was produced by exchanging the GFP in the St3Gal5-GFP vector for a Halo-tag amplified from the template pHTN-HaloTag (gift from Andrew Plested, Humboldt-Universität zu Berlin) using the restriction enzymes BamHI and HindIII.

### Primary rat hippocampal cultures and transfections

Primary hippocampal rat cultures were prepared as described previously (Kapitein et al., 2010) with minor modifications. Briefly, brains from E18/19 embryos or P0 pups were collected and the hippocampi were dissected and collected in HBSS (Thermo Fisher Scientific). After washing with HBSS, the hippocampi were incubated with trypsin (0.25%, Thermo Fisher Scientific) for 10 min at 37°C and then dissociated with a syringe (first 0.9 mm, then 0.45 mm). The dissociated cells were plated on poly-L-lysin (Sigma-Aldrich) coated coverslips at a density of 40.000-60.000 cells/mL in full medium (Dulbecco’s modified Eagle’s medium (DMEM; Gibco, Thermo Fisher), 10% fetal calf serum (FCS) (Gibco), 1× penicillin/streptomycin (Thermo Fisher Scientific), and 2 mM glutamine (Thermo Fisher Scientific). After 1 h the medium was exchanged to BrainPhys neuronal medium (STEMCELL Technologies) with SM1 supplement (STEMCELL Technologies) and 0.5 mM glutamine. Cells were incubated at 37°C, 5% CO2 and 95% humidity. Primary rat hippocampal cultures were transfected with Lipofectamin 2000 (Thermo Fisher Scientific) at a ratio of 1:2 DNA:Lipofectamin 2000 at DIV 12-16 according to the manufacturer’s instructions. Experiments were performed 24 to 48 hours after transfection. For the transfection, the conditioned medium was exchanged with BrainPhys (without supplements). After 1.5 h incubation with the transfection mixture, the medium was replaced again with the prewarmed conditioned medium.

### Immunocytochemistry (ICC)

For immunocytochemistry, cells were fixed with 4% Roti-Histofix (Carl Roth) / 4% sucrose for 10 minutes at room temperature (RT) and washed 3 times with PBS. The cells were then permeabilized (0.2%Triton X-100 (Supelco) in PBS) for 10 minutes, washed 2x with PBS and blocked with BBHE (BBHE: 0.1% TritonX, 10% heat-inactivated horse serum (Gibco), in PBS) for one hour at RT. When cells were transduced with St3gal5-GFP virus the signal was enhanced with an Atto488 pre-labeld GFP nanobody (Table S1). Primary antibodies were added in BBHE overnight at 4°C. Cells were washed again with PBS 3x before secondary antibodies were added in BBHE for 1 h at RT. Finally, the coverslips were washed 3 times with PBS and then mounted in mowiol (prepared according to the manufacturer’s instructions: 9.6 g mowiol 4-88 (Carl Roth), 24.0 g glycerin, 24 ml H_2_O, 48 ml 0.2 M Tris pH 8.5, including 2.5 g DABCO (Sigma-Aldrich)) on microscop slides before imaging.

### AAV production and transduction of primary hippocampal neurons

Adeno-associated viruses 9 (AAV9) were produced at the vector facility of the University Medical Center Hamburg-Eppendorf (UKE). Primary rat hippocampal cultures were transduced at DIV 16-17 with AAV9 carrying the synapsin-St3gal5-GFP vector (1 µl virus per 18 mm coverslip, viral genomes per µl= 4.6-5.6E+13 vg/µl) in conditioned medium. After 4-5 days, at DIV 20-21, the cells were fixed (see above) and used for further experiments.

### Cell lines and transfection of cell lines

HEK293T cells were cultured in full medium containing Dulbecco’s modified Eagle’s medium (DMEM; GIBCO, Thermo Fisher) and 10% fetal calf serum (FCS), 1× penicillin/streptomycin, and 2 mM glutamine at 37°C, 5% CO_2_, and 95% humidity. Cells were transfected with MaxPEI (Polysciences 24765-1) at 70% confluency. The DNA MaxPEI ratio was 1:3 and the transfection performed according to the manufacturer’s instructions. The cells were harvested 24-36 hours after transfection.

### Co-immunoprecipitation and immunoblotting

For co-immunoprecipitation, HEK293T cells grown in 10 cm dishes were transfected with a myosin VI dominant negative fused to EGFP or with EGFP alone as a control. Cells were harvested 24-36 hours after transfection. For the harvest, the cells were first washed with 4°C TBS (20 mM Tris HCl (pH 7.4), 150 mM NaCl, 0.1 % TritonX-100). Afterwards, cells from one 10 cm dish were lysed with 400 µl lysis buffer (50 mM HEPES pH 7.4, 300 mM NaCl, 0.5 % Triton X-100, complete protease inhibitor cocktail [Roche]). The lysate was centrifuged at 13.000 x g for 15 min and the supernatant (SN) was collected. At this stage, a sample of the SN was taken as “input”. The remainder of the SN was incubated with magnetic GFP-trap beads (Chromotek) on a rotor over night at 4°C. The beads were then washed 5x with lysis buffer (without protease inhibitor) using a magnetic rack before all lysis buffer was removed and the beads were resuspended in 80 µl 4x SDS buffer. The samples were then denatured at 98°C for 10 minutes and subjected to SDS-PAGE (10%) and immunoblotting. For the detection of myosin VI (full-length and dominant negative), the membrane was first developed with myosin VI antibody (Table S1) and in a second step incubated with GFP antibody (Abcam, Table S1) and developed a second time. The secondary antibody was ms-HRP coupled and detected with SuperSignal™ West Pico PLUS Chemiluminescent Substrate (Thermo Fisher Scientific).

### Pharmacological treatments

#### Latrunculin A (LatA) treatment

For the LatA treatment experiments, cells were incubated with a final concentration of 5 µM latrunculin A (2 mM stock in DMSO, Table S1) in prewarmed conditioned medium or aCSF (145 mM NaCl, 10 mM HEPES, 12,5 mM D-glucose, 1,25 mM NaH_2_PO_4_, 2,5 mM KCl, 1 mM MgCl_2_, 2 mM CaCl_2_, pH adjusted to 7,4) for 30 minutes at 37°C, 5% CO2 in 95% humidity. After 30 min incubation, the cells were imaged for a maximum of 45 minutes. In the control group, cells were treated with DMSO (Sigma-Aldrich).

#### Synaptotagmin antibody uptake assay

For the synaptotagmin antibody uptake assay, cells were incubated with pre-labeled synaptotagmin antibody (Table S1) (1:100) in aCSF for 30 minutes at 37°C, 5% CO2 in 95% humidity. After 30 min, the medium was exchanged to antibody-free medium for live imaging.

#### HaloTag labeling

When Halo-tag constructs were used, the conditioned medium was removed from the transfected cells and replaced with aCSF (see LatA treatment) containing the Halo-tag dye (Promega, according to manufacturer’s instructions, Table S1). After 30 minutes of incubation the medium containing the Halo-tag dye was replaced with the conditioned medium and cells were incubated again for at least 30 minutes before imaging.

### Wide-field, TIRF, and confocal spinning-disk microscopy

Live-cell microscopy and fixed cell imaging were performed with a Visitron Spinning-Disc TIRF-FRAP system on a Nikon Eclipse Ti-E controlled by VisiView software (Visitron Systems). Samples were kept in focus with the built-in Nikon perfect-focus system. The fluorophores were excited by 405, 488, 561 and 640 nm laser lines, coupled to the microscope via an optic fiber. The samples were imaged (confocal and TIRF) with a 100x TIRF oil objective (Nikon, ApoTIRF 100×/1.49 oil). Oblique and TIRF illuminations were obtained with an ILAS2 (Gattaca systems) TIRF system. Multichannel images were acquired sequentially using an appropriate filter set with an Orca flash 4.0LT CMOS camera (Hamamatsu). Confocal spinning disc imaging was performed with the Yokogawa CSU-X1 unit. For fixed cell imaging, multichannel z-stacks (distance 0.3 µm) were taken sequentially using an appropriate filter set with an Orca flash 4.0LT CMOS camera (Hamamatsu) or a pco.edge 4.2 bi sCMOS camera (Excelitas PCO GmbH). For live-cell imaging experiments, the images were acquired at 2 frames per second (single focal plane) or at specified indicated intervals. The samples were incubated in a stage top incubator (Okolab) at 37 °C, 5 % CO2 and 90 % humidity atmosphere. Neurons were imaged in regular culture medium unless specified otherwise. Coverslips were placed in a Ludin chamber (Life Imaging Services). If pharmacological treatment was performed during live imaging, the respective agent was added manually to the culture medium and incubated for the indicated time spans in the top stage incubator.

### Experimental design and statistical analyses

The experimental groups were blinded prior to data analysis.

#### Kymograph analysis

Kymographs were created with the Fiji plugin from KymographClear/ or KymographClear2a (Mangeol et al., 2016). For each cell, one axonal stretch was selected using the segmented line tool, and the line thickness was chosen to cover the axon thickness. GS trajectories shown in the kymograph were traced by hand using the straight-line tool. Moving trajectories (= slope) were traced separately from pausing/stationary events (= vertical lines). From the angle and length of the trajectories, the instant run length, the instant velocities and the instant pausing times were calculated. Times during which the GS did not move were divided into “pausing” (less than 60 s without processive movement) and “stationary” (more than 60 s without processive movement). To count pausing events and to localize them in the kymograph, the pointing tool was used to measure x/y values.

#### Analysis of pausing/stationary events at presynaptic boutons

To analyze pausing and stationary events at presynaptic boutons, axons were traced and boutons were selected according to their morphology with the FIJI segmented line tool in the timelapse image stack. The kymograph Distances between boutons were calculated using the Fiji macro MeasureSegementedDistances (https://doi.org/10.6084/m9.figshare.1133868.v1). Active boutons were distinguished from inactive boutons by live labeling with synaptotagmin antibodies. The distances between (active/inactive) boutons, measured in the timelapse videos were used to determine the position of (active/inactive) boutons (x value) along the corresponding kymograph. To calculate the distance of the pausing/stationary event (x value of the event) to the nearest bouton, the distance to all boutons was calculated and the smallest was selected. 1 µm distance to the next bouton was counted as pausing/being stationary at an (active/inactive) bouton. For an intrinsic control, random pausing events of the same number and in the same range to the corresponding kymograph were created in Excel. These random events were then treated in the same way as the “real” values to calculate the random pausing/being stationary % values. For statistics, a paired t-test was performed using Prism 9.1 (GraphPad).

#### GS counting

To count GS, primary hippocampal neurons were transduced or transfected with St3gal5 and fixed as described above. To distinguish the axon from the dendrite, ICC was performed and the AIS (ankyrin G) or presynaptic (bassoon) or somatodendritic compartment (MAP2) was labeled. The stained and mounted coverslips were imaged using a spinning disc confocal microscope. To enlarge the field of view, multiple images were taken using the scan slide tool and the resulting images were then stitched together using a Fiji macro. The images were then analyzed using Fiji. The axon was divided into three compartments: the proximal part (∼50-100 µm distance to the soma), the medial part (∼150-200 µm distance to the soma) and the distal part (at least ∼250 µm distance to the soma). The segmented line tool was used to measure the distance to the soma. Within each compartment, if possible, a stretch of axon was selected (proximal, medial: 40-72 µm; distal: 40-95 µm) and the GS vesicles were detected using the “Find Maxima” command with the threshold manually adjusted. The number of GS was then counted and normalized to 10 µm for each axonal area.

### Statistics

Statistical analyses were performed using Prism 9.1 (GraphPad), with detailed specifications of tests, significance levels, n numbers, and biological replicates included in the figure legends. Data are presented as individual dot plots, and error bars indicate means with their standard deviation across the manuscript. Statistical significance is defined as: n.s. for not significant, * for p < 0.05, ** for p < 0.01. For the kymograph analyses, run lengths and velocities are measured as the median per cell and presented as mean value for all cells. The pausing time is measured as average per cell and presented as mean value for all cells.

## Results

### GS are abundant in axons and are actively transported anterogradely and retrogradely

GS have been described and characterized previously in dendrites of hippocampal primary neurons ((Mikhaylova et al., 2016; Andres-Alonso et al., 2023). During earlier work, we noticed that vesicles positive for GS markers were also present in axons, which had not been reported in principal neurons before. We therefore set out to perform a detailed investigation of GS distribution, localization and trafficking in axons of rat hippocampal primary neurons. To visualize GS, we transduced or transfected dissociated primary neurons with the GFP-or Halo-tag coupled Golgi-targeting region of ST3 β-galactoside α-2,3-sialyltransferase 5 (St3Gal5), a tool previously described as a marker of GS and other Golgi-related compartments (Mikhaylova et al., 2016; Andres-Alonso et al., 2023). To identify neuronal subdomains, we performed immunocytochemistry (ICC) in adult neurons (DIV21-23) expressing St3Gal5 and stained for ankyrin G, an axon initial segment (AIS) marker, and MAP2, a somatodendritic marker. Indeed, confocal images showed that St3gal5 labels the somatic Golgi complex, dendritic Golgi outposts, GS in dendrites, as well as GS in the AIS, in the axon, and in the axonal growth cone (Fig. 1A). To further characterize the distribution of GS in the axon, we defined three areas: the proximal part (∼50-100 µm distance to the soma), the medial part (∼150-200 µm distance to the soma), and the distal part (at least ∼250 µm distance to the soma) (Figure 1B). We quantified the number of GS per 10 µm in these three areas and found that there are slightly more GS in the proximal part of the axon than in the medial or distal part (mean: proximal = 3.8 per 10 µm, medial = 3.1 per 10 µm, distal = 2.8 per 10 µm).

**Figure 1:**
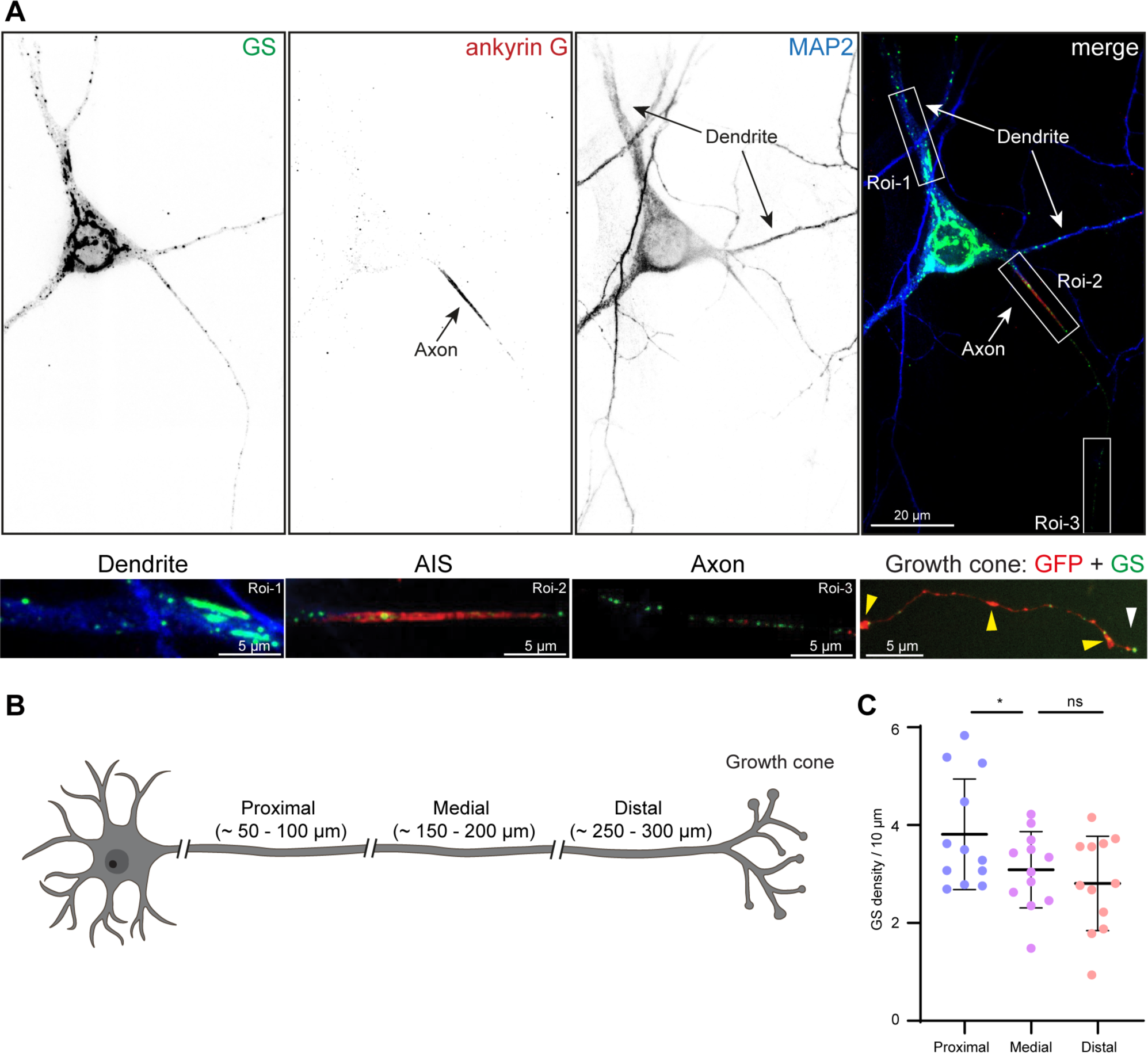
Distribution of GS in axons. A) Representative image of GS (green) in dendrites (Roi 1), in the AIS (Roi 2) and in the proximal axon (Roi 3) of primary hippocampal rat neurons (DIV16-17), stained with ankyrin G as an axon initial segment marker and MAP2 as a dendritic marker. GS were labeled with St3gal5-GFP and the signal intensity was boosted with an Atto488 prelabeled GFP nanobody. Right lower panel a representative image of an axonal neuronal growth cone transfected with St3gal5-HaloTag JF646 and eGFP, highlighting the boutons (yellow arrow) and the growth cone (white arrow) B) Schematic showing the axonal areas used for quantifying GS abundance in C. C) Quantification of GS distribution within the axon. Values are normalized to GS per 10 µm axon and show a significant difference for proximal area to the medial and no difference for the medial to the distal area (p* (proximal-medial) = 0.0405, p (medial-distal) = 0.4649; repeated measurements ANOVA test; n = 12). Values are presented as mean ± SD.

As previously described, GS are not only stationary but can be actively transported bidirectionally in dendrites (Mikhaylova et al., 2016). To investigate the mobility of GS in axons, we performed live-imaging of primary hippocampal neurons co-transfected with St3gal5-GFP to visualize GS and with actin-RFP, as it outlines the neuronal morphology and is enriched in dendritic spines, making it easy to identify the subcellular compartments (Figure 2A, video 1-2). To compare GS trafficking in axons and dendrites, we generated kymographs from time-lapse imaging stacks (Figure 2B). Similar to dendrites, we observed both immobile and bidirectionally moving GS in the axon. We decided to separately investigate four mobility parameters (van Bommel et al., 2019): being stationary (immobile for at least 60 s), pausing (immobile for less than 60 s), retrogradely transported or anterogradely transported. Interestingly, the stationary events (Figure 2C) occurred more frequently in the dendrites (mean = 2.4 per 10 µm) compared to the axon (mean = 1.5 per 10 µm), and pausing times were significantly longer in dendrites (mean = 18.8 s) than in axons (mean = 13.6 s). However, the number of pausing events per 10 µm, as well as the average run length and velocity was not significantly different between the compartments (Fig. 2D). Altogether, we found that GS are much more mobile in axons compared to dendrites (∼40 % vs. ∼ 15 % mobile, Figure 2E).

**Figure 2:**
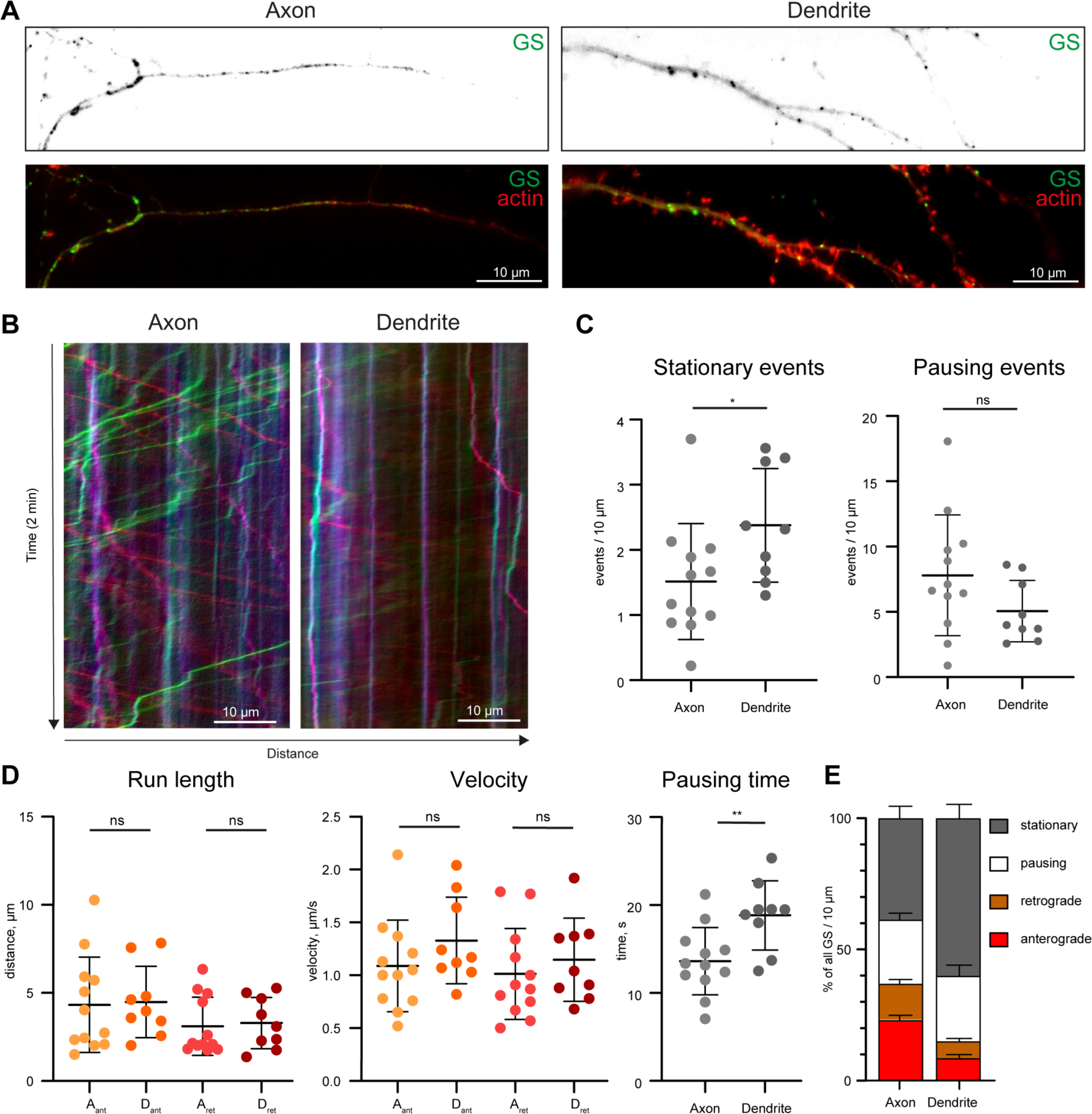
Trafficking of GS in the proximal axon. A) Representative confocal images of axons (left) and dendrites (right) of neurons expressing the GS-marker St3Gal5-GFP and actin-RFP. See also videos 1 (dendrite) and 2 (axon). B) Kymographs of the videos shown in A), imaged for 2 minutes at 4 frames per second (FPS), showing higher mobility of axonal GS compared to dendritic GS. C) Quantification of stationary and pausing events per 10 µm showing a significantly higher amount of stationary events (not moving for at least one minute) in dendrite than axons (p* = 0.0390; unpaired t-test; n(axon) = 12, n(dendrite) = 9; one axon or dendrite from one cell in 3 individual cultures;), whereas the pausing events (not moving for less than 60 s) were not significantly different (p = 0.1224; unpaired t-test), presented as mean ± SD. D) Quantification of GS trafficking parameters: run length, speed (divided in anterograde movement (Ant) and retrograde movement (Ret) and pausing time; same n as in C) in Axons (A) and in dendrites (D). The trafficking parameters are not significantly different (p (anterograde run length) = 0.8818, p (retrograde run length) = 0.7971, p (anterograde velocity) = 0.2135, p (retrograde velocity) = 0.4720; unpaired t-test) except for the pausing time, which is significantly longer in dendrites (p** = 0.0066; unpaired t-test). Values are presented as mean ± SD. E) Percental distribution of GS motility states in axons and dendrites: being stationary, pausing, being transported retrogradely or anterogradely displayed as % of the time the GS spend in that stage. Note a longer stationary time in dendrites as well as less retrograde and anterograde transport.

### GS pause at presynapses and are anchored to active boutons

As we observed both mobile and immobile GS in axons, we next asked whether GS pause or stop at specific axonal subcompartments. Since other organelles such as amphisomes and dense-core vesicles (DCV) are known to stall at presynaptic boutons (Andres-Alonso et al., 2019; Nassal et al., 2022), we asked whether immobile GS localize preferentially to presynaptic sites and whether this correlates with the synaptic activity at the bouton. To investigate this, we co-transfected primary hippocampal neurons with St3Gal5-GFP and with a cell fill (mRuby) to morphologically visualize presynaptic boutons. We then performed the synaptotagmin antibody uptake assay to distinguish between active and inactive boutons and took timelapse-images of live cells (Figure 3A, video 3). Images were taken in the medial/distal part of the axon because synapses are more frequently found in the distal part of the axon (Andres-Alonso et al., 2019; Qian et al., 2024). To analyze GS trafficking behavior, we created kymographs (Figure 3B) and selected the pausing and stationary events. We further investigated whether these events occurred at an active/inactive bouton (±1 µm) (Guedes-Dias et al., 2019). As an intrinsic control, we compared these results to a random distribution of events (pausing or stationary) along the kymograph (see Material & Methods). We found that short-term pausing events occurred significantly more often at boutons than would be expected from random pausing (Fig. 3C, left), regardless of the activity status of the bouton (Figure 3C, right). Interestingly, stationary GS were not only found preferentially at boutons (Fig. 3D, left), but were specifically enriched at active boutons (Figure 3D, right), suggesting that GS are actively recruited to active presynaptic sites.

**Figure 3:**
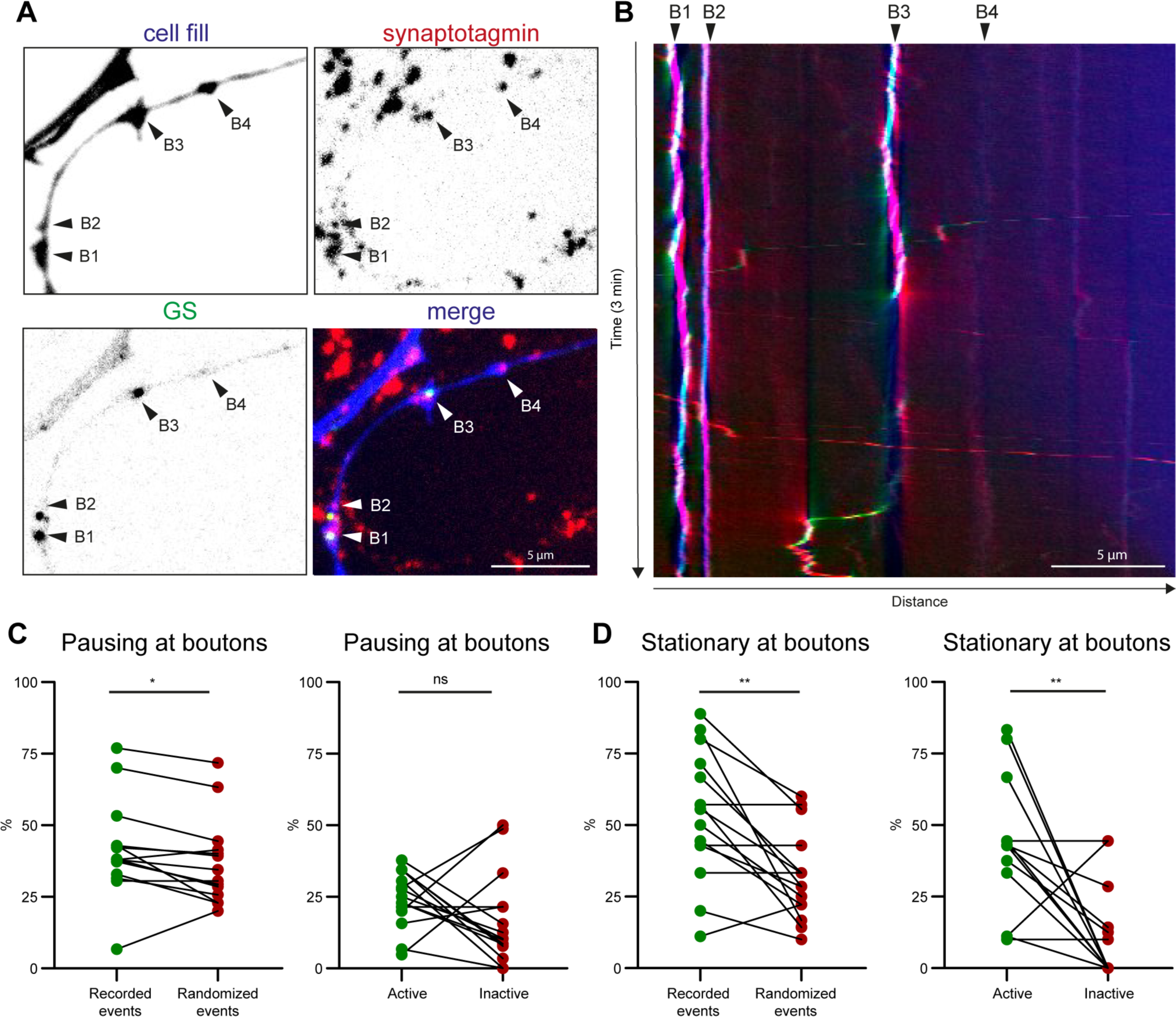
GS pause at presynaptic boutons and are anchored to active boutons. A) A representative image showing GS (green) at active presynaptic boutons (red) in DIV16-17 hippocampal neurons. Black arrows pointing to active boutons (B1-4); active boutons are labelled via synaptotagmin antibody uptake assay, also see video 3. B) Representative kymograph of the axon (shown in A) with red arrows pointing to the active boutons defined in A: (B1-4), showing multiple stationary events at active boutons. C) Description of GS pausing at boutons (left) but independent of bouton activity (right). For intrinsic control, GS pausing events are compared to random pausing events distributed along the examined axon (see Methods), (p* (GS-random) = 0.0199, p* (active-inactive) = 0.1716; paired t-test; n = 15, from three independent cultures) D) GS are stationary at boutons (left) and preferentially stationary at active boutons (right), same method and n as C) (p** (GS-random) = 0.0012, p** (active-inactive) = 0.0017; paired t-test).

### The axonal F-actin network disrupts processive trafficking of GS

The F-actin cytoskeleton is known to regulate the transport and localization of dendritic secretory organelles, such as lysosomes and the spine apparatus (van Bommel et al., 2019; Konietzny et al., 2022). It is also highly enriched in presynaptic boutons (Bingham et al., 2023). We therefore asked if presynaptic F-actin can regulate organelle positioning in distal axons. To determine the contribution of F-actin, we co-transfected primary hippocampal neurons with St3gal5-GFP and pORANGE-actin-TagRFP (CRISPR/Cas9-based endogenous actin knock-in label) to visualize GS and endogenous actin (Willems et al., 2020). Various F-actin structures exist in neurons, including a dense F-actin mesh in dendritic spines, F-actin patches in dendrites, longitudinal actin fibers, and a membrane periodic skeleton (MPS), consisting of F-actin, spectrin, and associated proteins in axons, dendrites, and the neck of dendritic spines, as well as F-actin patches at presynaptic sites (Xu et al., 2013; D’Este et al., 2015; Kevenaar and Hoogenraad, 2015; Bär et al., 2016; Konietzny et al., 2017). Local enrichments of F-actin are visible in confocal imaging as areas with a higher actin-TagRFP signal intensity (Fig. 4A, left). We then treated the cells with either the F-actin depolymerizing agent latrunculin A (LatA) (5 µM) or a solvent control (DMSO) for 30 minutes and performed time-lapse imaging (DIV16-17 video 4& 5) (Figure 4A). Upon LatA treatment, as expected, the actin-TagRFP labeling appeared smoother and more homogenous as the F-actin depolymerized (Fig. 4A, right). We created kymographs (Fig. 4B) to analyze GS trafficking and found a drastic effect on GS motility after depolymerization of F-actin following LatA treatment (Fig. 4C). The GS became faster (mean velocity: DMSO-Ant = 0.9 µm/s, LatA-Ant = 1.5 µm/s, DMSO-Ret = 0.8 µm/s, LatA-Ret = 1.3 µm/s), showed longer run lengths (mean run length: DMSO-Ant = 1.7 µm, LatA-Ant = 2.3 µm, DMSO-Ret = 1.5 µm, LatA-Ret = 2.1 µm) and shorter pausing times (mean DMSO = 10.8 s, mean LatA = 7.8 s). The number of stationary GS was not altered, suggesting that either stable LatA-insensitive F-actin structures or additional actin-independent mechanisms may also play a role in the long-term anchoring of GS. Altogether, these data demonstrate the important role of F-actin regulating GS trafficking in axons.

**Figure 4:**
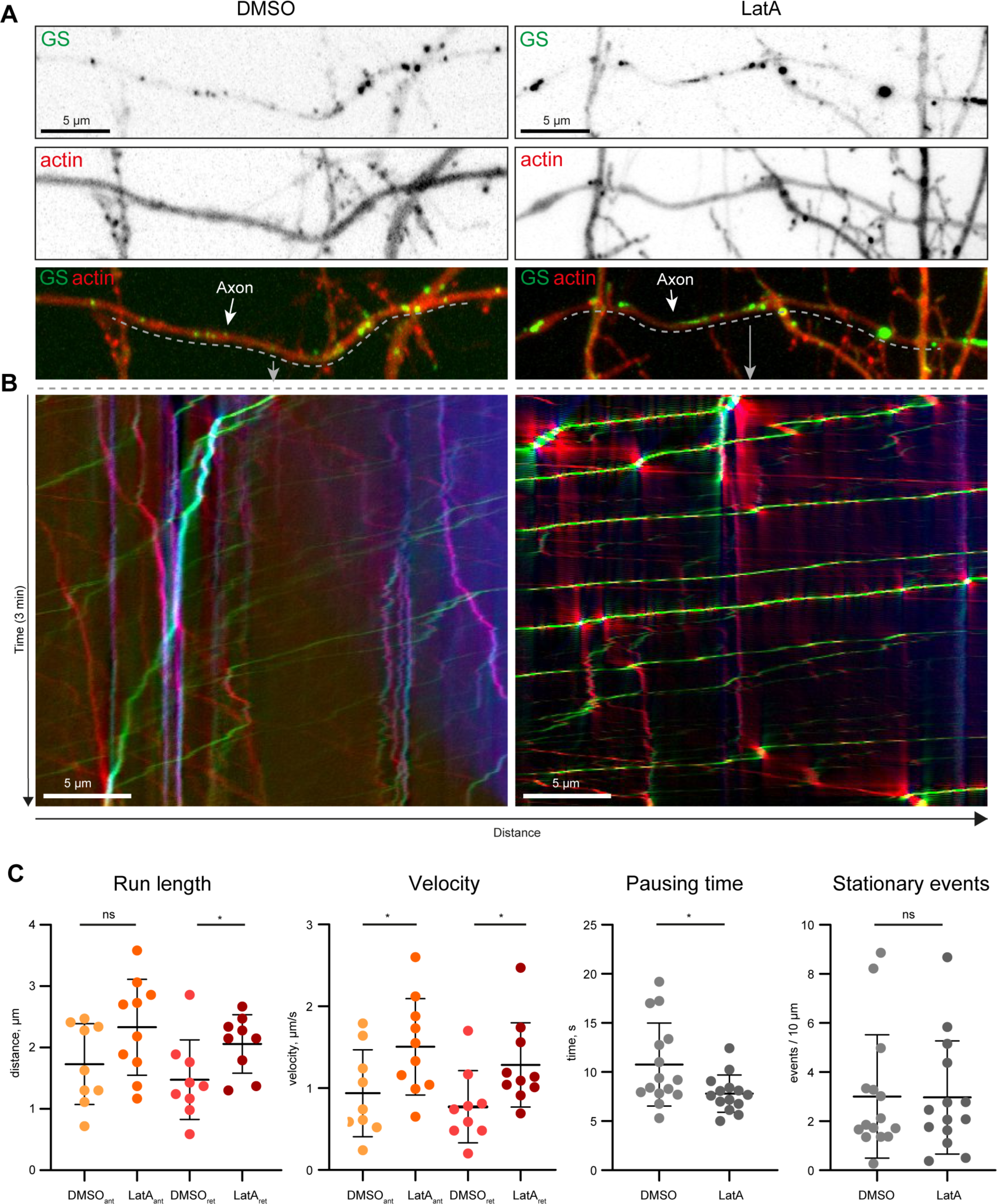
F-actin influences the trafficking of GS in axons. A) Representative confocal images of Golgi satellites in axons following latrunculin A (LatA) treatment (5 µM, right) and DMSO control treatment (left) in relation to actin (pORANGE-actin-tagRFP endogenous labelling and St3gal5-GFP overexpression). See also videos 4 (DMSO) and 5 (LatA). B) Kymographs of the videos shown in A), recorded for 3 min at 2 FPS, showing increased movement and decreased pausing of the GS after LatA treatment compared to DMSO ctr. C) Quantification of GS trafficking parameters: run length, velocity, pausing time and stationary GS; run length and velocity are increased in the LatA treatment group (p (anterograde run length) = 0.0891, p* (retrograde run length) = 0.0455, p* (anterograde velocity) = 0.0425, p* (retrograde velocity = 0.0334); unpaired t-test; n(DMSO) = 9, n(LatA) = 10; one axon from each cell in 3 individual cultures). The pausing time is significantly reduced by LatA treatment (p* = 0.0240; unpaired t-test; n(DMSO) = 15, n(LatA) = 14; one axon from each cell in 3 individual cultures). All values are presented as mean ± SD. Quantification of stationary events per 10 µm showing no difference in stationary events (p = 0.9659; unpaired t-test; n(axon) = 15, n(dendrite) = 14; one axon from one cell in 3 single cultures), presented as mean ± SD.

### Axonal GS are actively anchored to actin filaments by myosin VI motor protein

Long-term positioning of secretory organelles such as lysosomes at F-actin patches in dendrites is mediated by the motor protein myosin V (van Bommel et al., 2019). Myosin V is also known to stall dendritic cargo and reroute it into the dendrite (Janssen et al., 2017). Myosin VI, on the other hand, has been reported to be present in axons and to localize proteins to the axonal membrane (Lewis et al., 2011; Wagner et al., 2019). Furthermore, myosin VI can be activated by elevated Ca^2+^ levels induced by synaptic activity (Batters et al., 2016). We therefore asked whether myosin VI might be involved in the active localization of GS in the axon. To address this question, we used a GFP-tagged myosin VI dominant negative (MyoVI-dn) construct (Correia et al., 2008; van Bommel et al., 2019) in which the motor domain of the myosin VI heavy chain is removed, preventing it from binding to F-actin (Figure 5A). After overexpression, the construct competes with endogenous myosin VI for cargo binding. Myosin VI must dimerize to form a functional motor protein. Co-immunoprecipitation experiments indicated that MyoVI-dn can still interact with endogenous myosin VI (Figure 5B). This leads to the formation of a non-functional myosin VI (Figure 5A). To see if myosin VI is recruited to GS we co-transfected MyoVI-dn-GFP and St3gal5-Halo into primary hippocampal neurons and visualized the GS with the Halo dye JF646. Confocal imaging in medial and distal axons revealed colocalization of GS with MyoVI-dn-GFP, indicating that myosin VI is recruited to the surface of GS (Figure 5C). To investigate whether MyoVI-dn expression had an effect on GS trafficking and localization, we first quantified the density of GS in the medial/distal area of the axon and could show that MyoVI-dn increased the number of GS. Further, time-lapse confocal imaging followed by kymograph analysis (Figure 5D) revealed that although the pausing times of GS were not altered, the number of stationary GS was significantly reduced in the MyoVI-dn group (mean Ctr = 2.2 per 10 µm, mean MyoVI-dn = 1.3 per 10 µm) (Figure 5G). In conclusion, we found that there were overall more mobile GS, as the fraction of stationary GS was decreased in the MyoVI-dn group (Figure 5H). This suggests that myosin VI is important for the long-term anchoring of GS in the distal axon.

**Figure 5:**
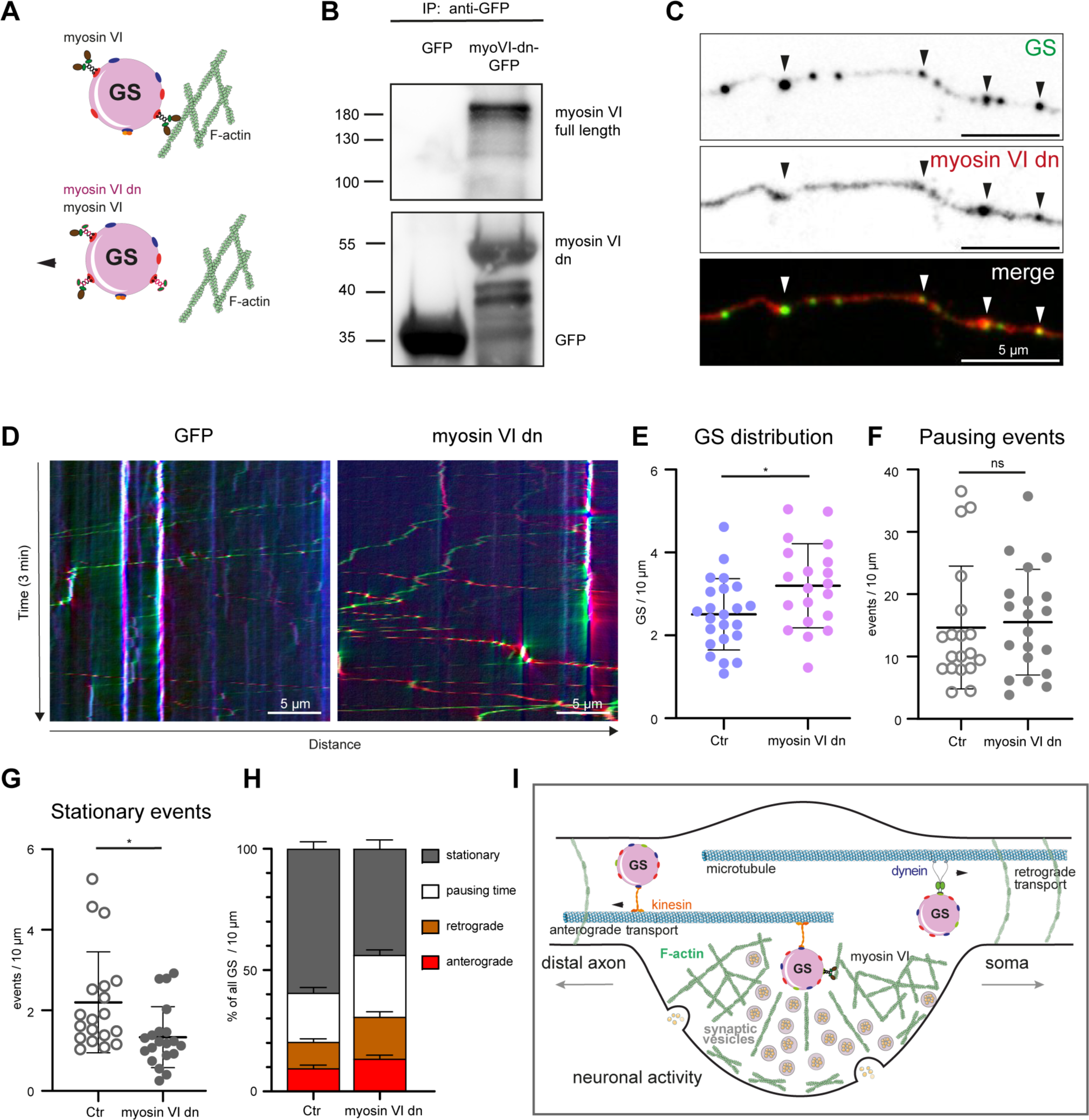
Myosin VI mediates the anchoring of GS in axons. A) Schematic of the myosin VI dominant negative (MyoVI-dn) principle. The motor domains of myosin VI are removed, preventing its interaction with actin. (This schematic was created with Servier medical Art templates, licensed under a Creative Commons Attribution 4.0 Unported Licencse; https://smart.servier.com) B) Western blot of MyoVI-dn-GFP co-IP from HEK293T cells extract compared to GFP ctr. showing interaction of MyoVI-dn with endogenous myosin detected with a myosin VI antibody (upper panel) and a GFP antibody (lower panel). C) Representative image showing the targeting of MyoVI-dn-GFP to GS (St3gal5-HaloTag-JF646). D) Representative kymographs of GS (St3gal5-HaloTag-JF646) in axons showing that overexpression of MyoVI-dn-GFP results in fewer stationary events compared to a GFP control. E) Quantification of the distribution of GS (St3gal5-HaloTag) in axons upon overexpression of MyoVI-dn-GFP compared to a control (GFP overexpression) showing a significant increase in GS per 10 µm upon MyoVI-dn overexpression in medial/distal axons (p* = 0.0206; unpaired t-test; n(Ctr) = 23, n(MyoVI-dn) = 20; one axon from one cell in 3 individual cultures), presented as mean ± SD. F) Quantification of pausing events per 10 µm. The pausing events are not altered (p = 0.7723; unpaired t-test; n(Ctr) = 19, n(MyoVI-dn) = 20; one axon/dendrite from one cell in 4 individual cultures), presented as mean ± SD. G) Stationary events show a significant reduction after MyoVI-dn overexpression (p* = 0.0126; unpaired t-test; same n as in F), presented as mean ± SD. H) Percent time of GS being stationary, pausing, retrogradely or anterogradely transported. Note a reduction in the pausing time in the MyoVI-dn group. I) Model showing the interplay of GS with F-actin and myosin VI at presynaptic boutons.

## Discussion

In this study, we show that Golgi satellites are present in the proximal, medial and distal parts of the axon as well as in the growth cone of primary hippocampal neurons. They are transported bidirectionally, frequently pause at boutons, and are preferentially recruited to active presynaptic sites. Furthermore, we show that myosin VI is involved in the long-term stalling of GS at active synapses, providing a mechanism for activity-dependent organelle localization.

We characterized the distribution of GS in the axon and found that GS are more abundant in the proximal part of the axon (50-100 µm from the soma) than in the more distal area. Overall, the trafficking behavior of GS in axons was similar to that in dendrites in terms of velocity and directionality of movement (anterograde vs. retrograde transport), while the fraction of stationary GS was higher in dendrites than in axons, and axonal GS were overall more mobile. In contrast to dendrites, microtubules in the axon have a uniform orientation with a plus end pointing to the periphery. Since we observed anterograde and retrograde transport of GS, we conclude that both plus-end and minus-end directed motors (i.e. kinesins and dynein) are involved in long-distance GS trafficking.

It is tempting to speculate that, similarly to dendrites, GS in axons constitute specialized microsecretory stations for posttranslational modification of secreted and transmembrane proteins, as well as local delivery of proteins and membrane components. Those functions involve direct interaction with many other organelles in the secretory pathway, such as ERGIC and endocytic vesicles, which makes it crucial that GS moving along the axon are stalled and recruited at the correct locations to enable such interactions. To investigate at which specific locations GS are stopped along the axon, we analyzed the stalling behavior of GS and could show that GS often pause for short time at presynaptic boutons and preferentially anchor for longer timespans at active synapses. The recruitment of organelles to presynaptic boutons upon neuronal activity has also been shown for other vesicles such as DCV or amphisomes, suggesting the existence of a general control mechanism of organelle positioning at presynapses.

How can active synapses recruit passing organelles? Guedes-Dias et al. have reported that dynamic MT ends at *en passant* synaptic boutons cause synaptic vesicles transported by the kinesin-3 motor protein to hop off the MT track (Guedes-Dias et al., 2019). However, presynaptic sites are also enriched in F-actin (Bingham et al., 2023), and we hypothesized that an additional mechanism, namely F-actin acting as a mesh to stop passing organelles, might be at play. A similar process has also been described in dendrites, where F-actin hot spots found along the dendrite at the base of spines or excitatory shaft synapses stop and anchor passing lysosomes (van Bommel et al., 2019). To test whether the F-actin network similarly affects GS trafficking in axons, we depolymerized F-actin using LatA and observed a dramatic effect on GS motility: they became faster, ran longer distances and paused for shorter amounts of time, clearly indicating that the F-actin network endogenously functions to slow down GS transport (van Bommel et al., 2019). Interestingly however, depolymerization of F-actin did not affect the number of stationary GS in axons. Of note, some F-actin structures, such as MPS, have low turnover of actin and are more resistant to LatA treatment and it remains to be determent if they can serve as organelle docking sites(Abouelezz et al., 2019). It is also possible that an interplay between MT dynamics and the F-actin cytoskeleton might regulate the recruitment of vesicles to *en passant* synapses, and that a combination of F-actin dependent and independent mechanisms may be involved in the long-term anchoring GS to boutons.

We next asked whether the stalling of GS at F-actin might be actively regulated by a myosin motor protein, similar to a previously described mechanism involving myosin V (van Bommel et al., 2019). Myosin V and VI are processive myosins involved in short-range cargo trafficking and localization. Both myosin V and VI have calmodulin binding regions through which they are activated in a calcium-dependent manner (Sellers et al., 2008; Batters et al., 2016; Shen et al., 2016). Myosin V can actively stall lysosomes at F-actin patches in dendrites (van Bommel et al., 2019), anchor the spine apparatus in dendritic spines, but it also anchors mRNA and vesicles containing dendritic proteins such as GluR1 and Kv4.2 at the AIS to be rerouted towards the dendrite (Lewis et al., 2009; Balasanyan et al., 2014; Janssen et al., 2017; Konietzny et al., 2022). This is in contrast to myosin VI, which is involved in the delivery of proteins to the axonal surface (Lewis et al., 2009). We therefore investigated the role of myosin VI in the axonal trafficking of GS and found that decreasing the amount of active myosin VI led to an increase in mobile GS, which goes in line with myosin VI actively stalling GS.

We therefore suggest a mechanism by which GS are recruited to active boutons by myosin VI, which can be locally activated by Ca^2+^ influx upon synaptic activity. The same mechanism could be true for other secretory vesicles, enabling their interaction at sites of high protein turnover, such as active synapses. In addition, GS may fuse with each other or with other secretory organelles in a myosin VI-dependent manner, as the fusion of autophagosomes and lysosomes is known to be myosin VI-dependent in non-neuronal cells (Tumbarello et al., 2012). This would explain why we observe more GS in distal axons in the MyoVI-dn group.

Axonal GS may be involved in various processes, including protein processing and delivery to lysosomes and autophagosomes. Recently it has been shown that trans-Golgi carriers deliver cathepsins to maturing lysosomes in axons of cortical neurons (Lie et al., 2021). It is possible that GS could carry similar functions and their role in this process remains to be elucidated. Interestingly, the Golgi components described by González et al. in axons of DRG neurons accumulate at nodes of Ranvier and might contribute to the trafficking of voltage gated sodium channels (González et al., 2016). The nodes of Ranvier contain F-actin meshwork which is important for the clustering of proteins like ankyrin G and NrCAM (González et al., 2016; D’Este et al., 2017). Since polysialylation of other adhesion proteins (NCAM) has been shown to be GS dependent in distal dendrites, one can speculate that similar modifications could be mediated by local GS for axonal adhesion proteins. Altogether, the axonal presence and activity-dependent localization of GS at synaptic boutons provides new possibilities for understanding and interpretations of mechanisms involved in local synaptic proteostasis.

## Supporting information

Reagents and resources

## Acknowledgments

The authors would like to thank B. van Bommel (AG membrane Biochemistry, FU, Berlin) for the cloning of the St3gal5-GFP-plasmid and I. Braren (Vector facility, UKE, Hamburg) for the production of the ST3gal5-GFP-AAV9. We are thankful to J. Bär, D. Hacker and L. Mallis (AG Optobiology, HU, Berlin) for the preparation of the dissociated rat hippocampal cultures. We thank A. Plested (HU, Berlin) and T. Oertner (ZMNH, UKE, Hamburg) for providing the plasmids indicated in Materials and methods. We thank E. Weisheim for providing a Fiji macro to stich the scan slide images.

This work was funded by the Deutsche Forschungsgemeinschaft Emmy Noether Programme MI1923/1-2, the Excellence Strategy – EXC-2049–390688087 and the DFG FOR5228 RP4.

## Video legends

**Video 1: GS trafficking in axons of primary hippocampal neurons**

Representative video showing GS (St3gal5-GFP in green) and Actin-RFP (red) imaged with 4 Frames per second (FPS) for 2 minutes in dissociated primary rat hippocampal neurons (DIV 16-17).

**Video 2: GS trafficking in dendrites of primary hippocampal neurons**

Representative video showing (St3gal5-GFP in green) and Actin-RFP (red) imaged with 4 FPS for 2 minutes in dissociated primary rat hippocampal neurons (DIV 16-17).

**Video 3: GS pause at active presynaptic boutons**

Representative video showing (St3gal5-GFP in green), mRuby (blue) and active boutons (synaptotagmin antibody assay, red) imaged with 2 FPS for 3 minutes in dissociated primary rat hippocampal neurons (DIV 16-17).

**Video 4: GS trafficking in axons, solvent control (DMSO)**

Representative video showing GS (St3gal5-GFP in green) and endogenous actin (pORANGE-actin-TagRFP; CRISPR/Cas9-based endogenous actin knock-in label in red) imaged with 2 FPS for 3 minutes in dissociated primary rat hippocampal neurons (DIV 16-17).

**Video 5: GS trafficking in axons, latrunculin A treatment**

Representative video showing GS (St3gal5-GFP in green) and endogenous actin (pORANGE-actin-TagRFP; CRISPR/Cas9-based endogenous actin knock-in label in red) imaged with 2 FPS for 3 minutes in dissociated primary rat hippocampal neurons (DIV 16-17).

